# MR Corge: Sensitivity analysis of Mendelian randomization based on the core gene hypothesis for polygenic exposures

**DOI:** 10.1101/2024.07.18.604191

**Authors:** Wenmin Zhang, Chen-Yang Su, Satoshi Yoshiji, Tianyuan Lu

## Abstract

**Summary:** Mendelian randomization is being utilized to assess causal effects of polygenic exposures, where many genetic instruments are subject to horizontal pleiotropy. Existing methods for detecting and correcting for horizontal pleiotropy have important assumptions that may not be fulfilled. Built upon the core gene hypothesis, we developed MR Corge for performing sensitivity analysis of Mendelian randomization. MR Corge identifies a small number of putative core instruments that are more likely to affect genes with a direct biological role in an exposure and obtains causal effect estimates based on these instruments, thereby reducing the risk of horizontal pleiotropy. Using positive and negative controls, we demonstrated that MR Corge estimates aligned with established biomedical knowledge and the results of randomized controlled trials. MR Corge may be widely applied to investigate polygenic exposure-outcome relationships.

**Availability and Implementation:** An open-sourced R package is available at https://github.com/zhwm/MRCorge.

## Introduction

Mendelian randomization (MR) is a promising tool for assessing potential causal relationships^1,2^ since randomized controlled trials are often limited by ethical and practical constraints. Using genetic variants as instrumental variables, MR may effectively mitigate the risks of confounding and reverse causation that frequently bias observational association studies^1,2^. MR relies on essential instrumental variable assumptions of relevance, independence, and exclusion restriction^3,4^. Specifically, genetic instruments must be strongly associated with the exposure, be independent of any confounders, and affect the outcome only through the exposure. The relevance assumption can be fulfilled by selecting genetic variants demonstrating genome-wide significance in genome-wide association studies (GWAS) of exposures, while the independence assumption can be examined by testing for instrument-confounder associations. However, exclusion restriction, also known as the assumption of no horizontal pleiotropy, remains challenging to verify.

To robustly derive MR estimates in the presence of invalid genetic instruments, several methods have been developed beyond the original Wald ratio and inverse variance weighted (IVW) methods^5^. For instance, the median-based methods (e.g. simple median, weighted median, and penalized weighted median methods) require that at least 50% of the instruments or their weights are valid^6,7^. The mode-based methods (e.g. simple mode and weighted mode methods) require that the mode of the causal effect estimates or their weights is based on valid instruments^8^. The MR PRESSO method identifies and removes outlier instruments^9^, while the contamination mixture method identifies one or more groups of instruments with consistent causal estimates^10^, both having the assumption that a plurality of the instruments should be valid^9,10^. The MR Egger method and MR RAPS method allow for pervasive pleiotropic effects but have an additional assumption that the strength of the genetic instruments is independent of their direct effects on the outcome (i.e. InSIDE assumption)^11,12^.

In recent years, MR has been increasingly utilized to estimate the potential causal effects of polygenic exposures, for which tens or hundreds of independent genetic variants can be identified as candidate instruments from GWAS of rapidly growing sample sizes^13,14^. It has been recognized that many, sometimes the vast majority, of these candidate instruments do not have a biologically interpretable role in the exposures and can be involved in multiple distinct biological pathways^15–17^. This substantially increases the risk of violating the additional assumptions of the aforementioned MR methods.

The core gene hypothesis in the theory of the omnigenic model^15^ has been proposed to explain the high polygenicity of complex traits. The core gene hypothesis states that genetic variants within core genes or pathways have more direct and specific effects on a polygenic trait than those within peripheral genes or pathways, which more likely play regulatory roles. We hypothesize that for polygenic exposures, using instruments from core genes or pathways may reduce the risk of horizontal pleiotropy.

In this work, we present MR Corge for performing sensitivity analysis of MR based on the core gene hypothesis. MR Corge estimates the effect of a polygenic exposure using a group of putative core instruments. We demonstrate the utility of MR Corge in assessing polygenic exposure-outcome relationships using positive and negative controls.

## Methods

MR Corge relies on two important assumptions derived from the core gene hypothesis^15^. First, for a polygenic exposure, a small number of core genes have a direct biological impact on the trait. Second, genetic instruments with a stronger variant-trait association (i.e. core instruments) are more likely to act through core genes, whereas those with a weaker variant-trait association (i.e. peripheral instruments) are more likely to act through peripheral genes and be subject to horizontal pleiotropy. With these assumptions, MR estimates obtained based on the putative core instruments are less likely to be biased by horizontal pleiotropy than those based on all instruments.

In MR Corge, the most crucial step is to identify putative core instruments from all candidate instruments. Four instrument ranking options have been made available, based on

1. 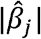 (default setting): the per-allele effect of the *j*-th genetic variant on the exposure, estimated in the GWAS of the exposure;
2. 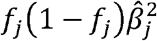 : the per-variant heritability of the *j* -th genetic variant, which is proportional to the squared per-allele effect normalized by the minor allele frequency (*f*_*j*_);
3. 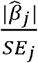: the significance of the *j*-th genetic variant in the GWAS of the exposure;
4. 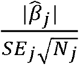: the sample size (*Nj*)-normalized significance of the *j*-th genetic variant, which can differ from (3) especially in meta-analyses of GWAS with multiple participating studies.

All instruments are ranked based on one of these four metrics in descending order. Among the tied instruments, ranks are randomly assigned. Then, the instruments are categorized into *K* groups, where each group contains approximately the same number of instruments. In practice, we recommend choosing *K* such that each group contains 5 to 10 instruments.

MR Corge then generates group-specific and cumulative MR estimates that can be based on any existing methods (Figure 1A). By default, MR Corge applies the IVW, weighted median, and weighted mode methods for estimation. The *K*-th cumulative MR estimate (*k* ∈ {1, …, *K*}) is obtained based on all instruments in the *K*-th and all preceding groups, if any. An F-statistic is calculated for each group of instruments to assess whether the MR estimates may be subject to weak instrument bias^18^. MR estimates based on the first group of instruments, which are putative core instruments, are the primary MR Corge estimates.

**Figure 1.**
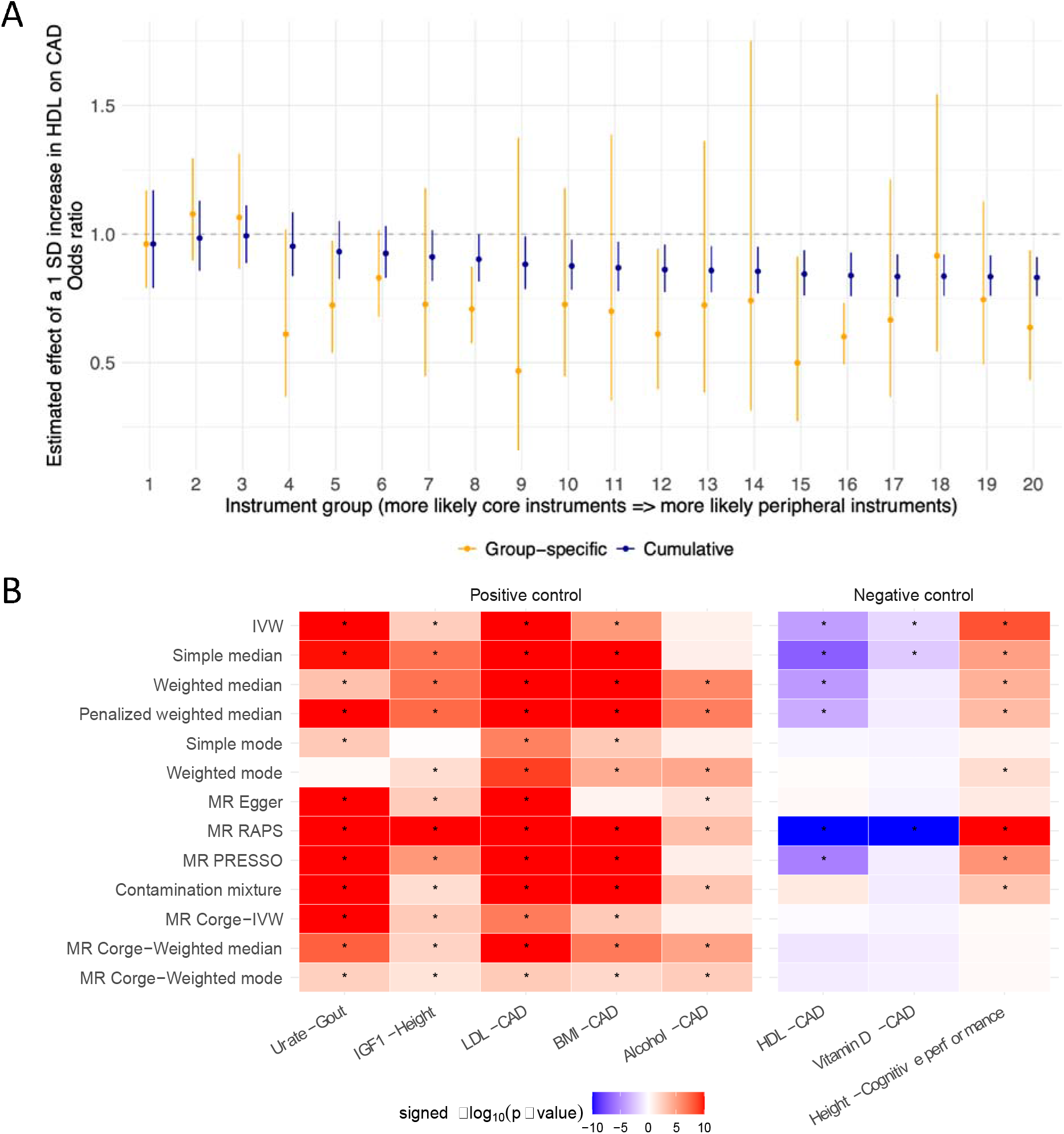
Benchmarking the performance of MR Corge. (A) Group-specific and cumulative MR Corge estimates of the effect of serum high-density lipoprotein cholesterol (HDL) levels on the risk of coronary artery disease (CAD). Estimates obtained using the inverse variance weighted method are illustrated with error bars indicating 95% confidence intervals. SD, standard deviation. (B) Significance of association estimates in positive and negative controls. Asterisks (*) indicate estimates with a p-value < 0.05. P-values <1.0×10^−10^ were set to 1.0×10^−10^ for visualization purposes. Red colors represent that increased exposure was predicted to increase the outcome while blue colors represent that increased exposure was predicted to decrease the outcome. Instruments were ranked based on the default ranking criterion based on the absolute value of per-allele effect size.

## Applications

We utilized several positive and negative controls to examine whether sensible MR estimates could be obtained using MR Corge.

The positive controls are

1. Serum insulin-like growth factor 1 (IGF1)-height association: IGF1 mediates the effects of growth hormone in various tissues, promotes the proliferation and differentiation of chondrocytes and myoblasts, and supports energy production, resulting in bone elongation and height increase^19–21^;
2. Serum urate-gout association: Gout is primarily caused by elevated levels of circulating urate, which can form urate crystals in joints and trigger inflammatory responses^22^;
3. Serum low-density lipoprotein cholesterol (LDL)-coronary artery disease (CAD), body mass index (BMI)-CAD, and alcohol intake-CAD associations: The risk-increasing effects of hyperlipidemia, obesity, and excessive alcohol intake on CAD have been confirmed in previous randomized controlled trials as well as Mendelian randomization and observational studies^23–27^.

The negative controls are

1. Serum high-density lipoprotein cholesterol (HDL)-CAD and vitamin D-CAD associations: Although increased HDL and vitamin D levels have been associated with a decreased risk of CAD in observational studies, randomized controlled trials have shown no meaningful protective effects of HDL-increasing drugs or vitamin D supplements against CAD in the general population with normal HDL or vitamin D levels^28–31^;
2. Height-cognitive performance association: Height is a significant predictor of cognitive performance in observational studies^32^, yet this association can be strongly confounded by various factors, such as socioeconomic status, nutrition, and assortative mating. We assessed the height-cognitive performance association using LD score regression (version 1.0.1)^33,34^ to estimate the genetic correlation based on within-sibship studies, which can effectively control for unmeasured confounders^35^.

GWAS summary statistics of these exposures and outcomes were obtained from large-scale studies (**Supplementary Table S1**)^36–44^. For each exposure, we performed linkage disequilibrium (LD) clumping using the ieugwasr R package (version 1.0.1)^45^ with an LD reference panel generated from 50,000 randomly sampled UK Biobank participants of European ancestry^46^. We retained independent genetic variants (LD r^2^ < 0.001, window size = 1 Mb) that demonstrated genome-wide significance (p-value < 5.0×10^−8^) as candidate instruments. The summary statistics of these instruments were obtained from the GWAS of the corresponding outcomes. When an instrument was not present, we attempted to identify its proxy variant with an LD r^2^ > 0.8 using the same LD reference panel. Data harmonization was performed using the TwoSampleMR R package (version 0.5.7)^45^. Palindromic variants a minor allele frequency > 0.42 were excluded. MR Corge estimates were obtained using the IVW, weighted median, and weighted mode methods, with *K*= 5 for the alcohol-CAD association, *K*= 20 for the IGF1-height, urate-gout, LDL-CAD, BMI-CAD, HDL-CAD, and vitamin D-CAD associations, and *K* = 100 for the height-cognitive performance association, where each instrument group contains 5 to 10 instruments. All genetic instruments and their functional impacts on nearby genes annotated using the Ensembl Variant Effect Predictor (release 112)^47^ are detailed in **Supplementary Table S2**.

We also derived MR estimates for each of these exposure-outcome pairs based on all instruments, using the IVW, simple median, weighted median, simple mode, weighted mode, penalized weighted median, MR Egger, MR RAPS, MR PRESSO, and contamination mixture methods. All MR estimates are provided in **Supplementary Tables S3** and **S4**.

## Results

### Positive controls

Using the default instrument ranking criterion based on the absolute value of per-allele effect size, the putative core instrument groups contained genetic variants mapping to genes with well-established biological functions related to the exposures. For instance, the putative core instruments for serum IGF1 levels included rs856540, an intergenic variant near IGFBP3 (main carrier of IGF1), while the putative core instruments for serum urate levels included rs1165196, a missense variant of SLC17A1, and rs1171614, an intronic variant of SLC16A9, where both SLC17A1 and SLC16A9 are important urate solute carriers. MR Corge estimates consistently showed that increased serum IGF1 levels could lead to increased height while increased serum urate levels could elevate the risk of gout (**Figure 1B**). Meanwhile, serum LDL levels and BMI were predicted to increase the risk of CAD (**Figure 1B**). Without selecting instruments, the simple mode, weighted mode, and MR Egger methods did not identify all of these associations (**Figure 1B**).

The strongest instrument for alcohol intake was rs1229984, a missense variant of ADH1B (alcohol dehydrogenase). ADH1B regulates the accumulation of acetaldehyde which can cause unpleasant effects such as flushing, nausea, and rapid heart rate^48,49^. However, most of the other instruments do not have a direct role in the metabolism of alcohol. With the core instruments, MR Corge estimates obtained using the weighted median and weighted mode methods identified the risk-increasing effect of alcohol intake on CAD, although the MR Corge estimate using the IVW method, along with four other methods based on all instruments, did not identify this effect (**Figure 1B**).

### Negative controls

Furthermore, MR Corge estimates consistently showed that serum HDL and vitamin D levels may not be causally associated with CAD, while based on all instruments, the IVW, simple median, weighted median, penalized weighted median, MR RAPS, and MR PRESSO suggested protective effects of at least one exposure (**Figure 1B**). The putative core instruments for HDL levels included rs9989419, an intergenic variant near CETP (cholesteryl ester transfer protein, transferring cholesterol esters from HDL particles to other lipoproteins^50^), while the putative core instruments for vitamin D levels included rs117576073, an intronic variant of CYP2R1 (vitamin D 25-hydroxylase), rs4694423, an intergenic variant near GC (vitamin D binding protein), and rs12803256, an intergenic variant near DHCR7 (7-dehydrocholesterol reductase which balances biosynthesis of cholesterol and vitamin D^51^). Interestingly, consistent with MR Corge estimates, CETP-inhibitors have been a recent focus of drug development yet have not shown effectiveness in reducing cardiovascular disease risks in randomized controlled trials, despite their capacity to significantly elevate serum HDL levels^28,29^.

Additionally, MR Corge estimates indicated that height was not associated with cognitive performance, consistent with the absence of genetic correlation based on within-sibship studies (Supplementary Table S5). In contrast, when all instruments were utilized, eight of the ten MR methods suggested a causal effect of increased height on increased cognitive performance (Figure 1B).

## Discussion

Instrumental variable assumptions are essential for deriving valid MR estimates. It has been recognized that carefully selecting genetic instruments which represent relevant biological mechanisms may mitigate the risk of horizontal pleiotropy, such as using lead variants in cis-quantitative trait loci for instrumenting the abundances of mRNA transcripts, proteins, and certain metabolites^52–57^. However, assessing potential causal effects of many polygenic exposures using MR remains challenging since the underlying biological mechanisms may not have been fully elucidated and a large number of candidate instruments can be subject to horizontal pleiotropy. MR Corge can be a useful tool for performing sensitivity analysis for such exposures.

With positive and negative controls, we demonstrated that MR estimates based on all instruments could be inconsistent with established biomedical knowledge or the results of randomized controlled trials, even when robust MR methods were used. This inconsistency is likely due to the violation of additional assumptions of these methods. In contrast, with a small number of putative core instruments, the assumptions of robust MR methods, such as the weighted median and weighted mode methods, are more likely to be fulfilled due to the enrichment of valid instruments. As a result, MR Corge could consistently provide sensible MR estimates. Moreover, because the putative core instruments had the strongest associations with the exposures, the F-statistics of the core instrument groups were higher than the overall F-statistics based on all instruments (**Supplementary Table S4**), indicating a low risk of introducing weak instrument bias^18^ and further supporting that appropriate instrument selection should prevail over maximizing the proportion of exposure variance explained by the instruments. This principle likely also applies to analyses using genetic risk scores as instrumental variables, where existing methods for addressing horizontal pleiotropy are not applicable.

Notably, a universal rule for determining core instruments may not exist, and the core instrument-peripheral instrument classification may not be binary^15^. Empirically, we found that ranking based on the absolute value of per-allele effect size outperformed other ranking criteria (Supplementary Figure S1). However, interpreting MR Corge estimates requires cautious investigation of the putative core instruments, as well as triangulation of multiple lines of evidence, as with other MR or epidemiological methods.

In summary, built upon the core gene hypothesis, MR Corge may identify putative core instruments to obtain MR estimates that are less likely to be biased by horizontal pleiotropy. MR Corge may have wide utility as a tool for performing sensitivity analysis when the exposures demonstrate high polygenicity, while the underlying biological mechanisms remain unclear.

**Supplementary Figure S1.**
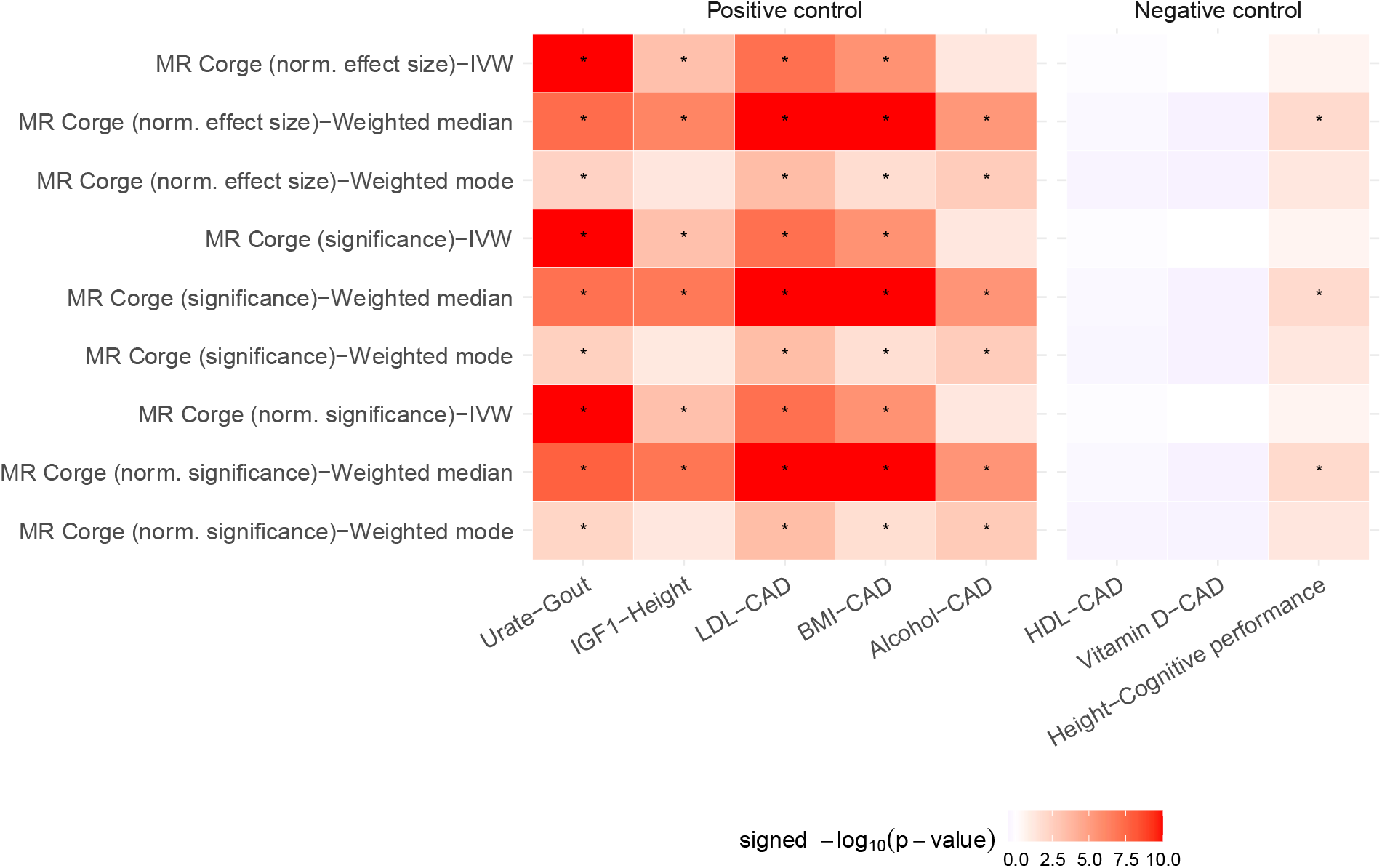
Significance of MR Corge estimates in positive and negative controls using alternative instrument ranking criteria. Asterisks (*) indicate estimates with a p-value < 0.05. Red colors represent that increased exposure was predicted to increase the outcome while blue colors represent that increased exposure was predicted to decrease the outcome. P-values <1.0×10^−10^ were set to 1.0×10^−10^ for visualization purposes.

